# Divergent Subregional Information Processing in Mouse Prefrontal Cortex During Working Memory

**DOI:** 10.1101/2024.04.25.591167

**Authors:** Alex Sonneborn, Lowell Bartlett, Randall J. Olson, Russell Milton, Atheir I. Abbas

## Abstract

Working memory (WM) is a critical cognitive function allowing recent information to be temporarily held in mind to inform future action. This process depends on coordination between key subregions in prefrontal cortex (PFC) and other connected brain areas. However, few studies have examined the degree of functional specialization between these subregions throughout the phases of WM using electrophysiological recordings in freely-moving animals, particularly mice. To this end, we recorded single-units in three neighboring medial PFC (mPFC) subregions in mouse – supplementary motor area (MOs), dorsomedial PFC (dmPFC), and ventromedial (vmPFC) – during a freely-behaving non-match-to-position WM task. We found divergent patterns of task-related activity across the phases of WM. The MOs is most active around task phase transitions and encodes the starting sample location most selectively. Dorsomedial PFC contains a more stable population code, including persistent sample-location-specific firing during a five second delay period. Finally, the vmPFC responds most strongly to reward-related information during the choice phase. Our results reveal anatomically and temporally segregated computation of WM task information in mPFC and motivate more precise consideration of the dynamic neural activity required for WM.

## Introduction

Working memory (WM) is a fundamental cognitive function allowing prior sensorimotor and rule information to be held in mind, manipulated, and protected from interference for future use^1^. This process relies on a distributed hierarchy of brain networks performing varying degrees of top-down and bottom-up operations^2,3^. The prefrontal cortex (PFC) is positioned at the top of this hierarchy, exerting the highest-order influence over WM via extensive reciprocal connections with cortical^4^ and subcortical structures^5^. It is thought to be responsible for orchestrating several key aspects of WM, including actively directing and maintaining attention toward salient features of a context, selecting strategies to accomplish goals based on contextual needs, and monitoring the outcome of enacted motor plans to change strategies if necessary^6,7^. Difficulty with any of these functions is a prevalent symptom across many human neurological and psychiatric disorders, and usually coincides with aberrant activation of PFC-containing networks during WM^8,9^. A deeper examination of PFC activity throughout the different phases of WM is critical to better understanding the potential mechanisms underlying diverse types of WM dysfunction.

Over the past decade, mice have become a standard model organism in PFC research due to rapid development of genetic tools permitting more precise targeting of cell-types and brain-wide circuits^10^. Extensive connectomic and genomic mapping has established that mouse PFC can be divided into subregions based on local and long-range projection patterns^11,12^ and cytoarchitecture^13,14^. However, attempts at segregating mouse PFC into subregions based on functional processing of WM task features has yielded surprisingly inconsistent findings^15^. A potential explanation for this variability comes from recent work showing that neural activity subserving goal-directed actions is spread across many brain areas, and the primary locus of control can shift dynamically depending on contextual needs and temporal progress through a task^16–22^. Thus, trying to localize multi-faceted mental processes, like WM, onto isolated mouse PFC subregions may be an unreliable approach. Instead, experiments in mice should focus on characterizing a range of WM-related computations in multiple PFC subregions over the entire time course of a single behavioral paradigm^15^.

Most mouse studies probing PFC neural circuit contributions to spatial WM have concentrated on single subregions within the same task^23–26^, and reports using electrophysiological recordings from multiple subregions are not accompanied by a detailed comparison between them^19,27–29^. Moreover, most multi-regional WM data have been collected from head-fixed mice^30–33^. Critically, more comprehensive analysis describing how distinct PFC areas selectively contribute to WM task variables across time in freely-moving mice is needed, as head-fixed experiments may engage different brain networks than more naturalistic behaviors^34,35^. To this end, we recorded single-units in three adjacent mouse PFC subregions agreeing with modern PFC parcellation schemes^15^: the supplementary motor cortex (MOs), the dorsomedial PFC (dmPFC), and the ventromedial PFC (vmPFC). Activity in these subpopulations was tracked in real time as the mice performed a freely-behaving delayed-non-match-to-position WM task ^36^. We asked how each subregion represented the retrospective sample location and other task-related information such as prospective choices and reward across the encoding, maintenance, retrieval, and outcome phases of WM^37^. Our results indicate that the dmPFC population stably codes for the retrospective sample location throughout all phases in single behavioral trials, while MOs prioritizes contextual transitions and vmPFC is most sensitive to choice and outcome variables.

## Results

### Behavior and electrode implantation

To examine the relationship between spatial WM and PFC subregional activity, mice were water restricted to ∼90% of their initial body weight and trained on a freely-moving, delayed-non-match-to-position (DNMTP) WM task (Fig. 1*A*). Critically, this task was designed so that mice could not know the exact choice port they would need to visit until the end of the delay period. During training, mice that made the correct non-match choice on ≥70% of trials over three consecutive days were implanted with a custom-built, 28-wire, advanceable bundle of microelectrodes into one of three separate PFC subregions (see **Methods** for details): the supplementary motor area (MOs, blue, n = 4), dorsomedial PFC (dmPFC, green n = 6), or ventromedial PFC (vmPFC, pink, n = 6) (Fig. 1*B*). We gathered at least three daily sessions of simultaneous behavioral and electrophysiological data per mouse, advancing the electrodes ventrally into the PFC by ∼60 µm after each session, so that new neurons were recorded the following day (white dots in Fig. 1*B* depict the final electrode bundle locations for each mouse). Single-units were isolated offline using Kilosort3, aligned to important DNMTP task events, and organized into pseudopopulations by combining the neurons recorded over all sessions within each subregion. The total neuron counts for each pseudopopulation were 304 in the MOs, 354 in the dmPFC, and 330 in the vmPFC. A one-way ANOVA revealed no significant main effect of implant location on DNMTP performance (F(2,70) = 2.71, p = 0.073, Fig. 1*C*).

**Figure 1.**
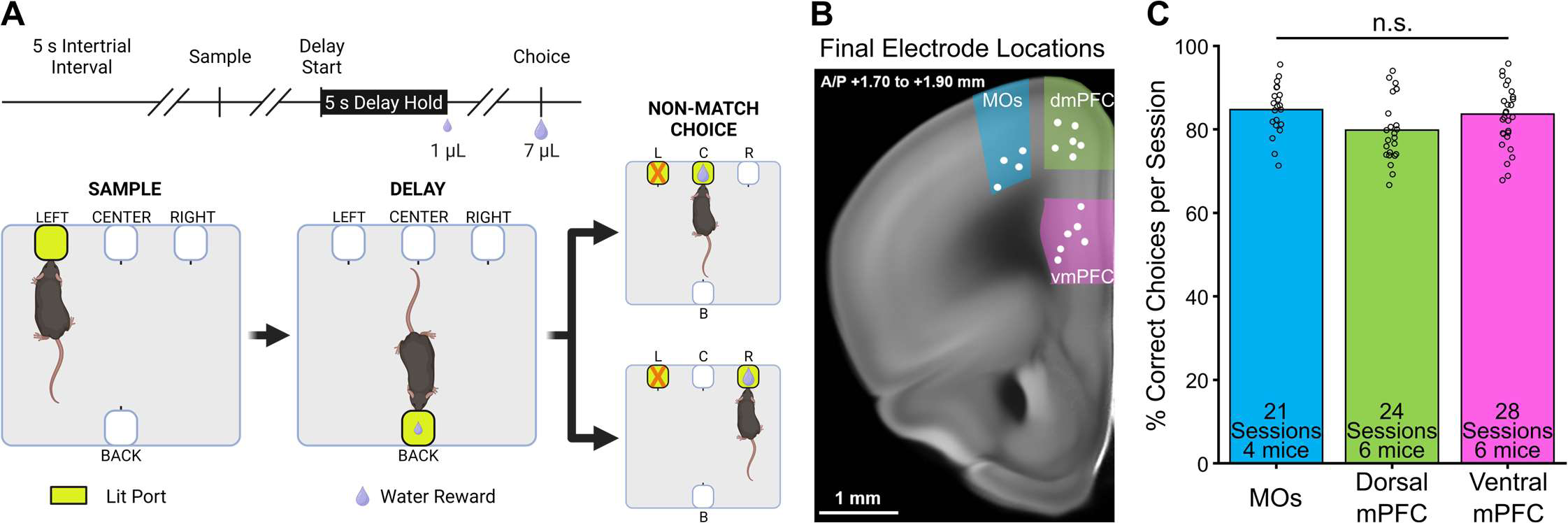
A freely-moving delayed non-match-to-position task allows for the examination of prefrontal neural activity during spatial working memory in mice. ***A,*** Schematic of the delayed non-match-to-position task. **(Top)** Time course of a single trial. Diagonal parallel slashes represent variable amounts of time between task phases. **(Bottom)** Diagram of a correct trial. Progression through the task starts with the sample phase (*left panel*), during which one of the two outer front ports lights up. The lit sample port location has a 50% chance of being on either the left or right side, but only a left side example trial is shown here. The mouse pokes its nose into the lit sample port to make the delay port on the opposite side (“back”) of the box available (*center panel*). Then the mouse pokes and holds its nose in the delay (“back”) port for five seconds, after which it receives a 1 µL reward. This leads to a choice phase (*right panel*), where both the initial sample port and one of the remaining other front ports lights up, and the mouse is required to nose poke the lit port that it has not visited previously. Non-match choices also have a 50% chance of being either of the two non-sample locations on any given trial. A much larger 7 µL reward is dispensed after a correct non-match choice. Panel A was created with BioRender.com. ***B,*** Mice were implanted with recording electrodes in either MOs (blue), dorsal mPFC (green), or ventral mPFC (pink). Example coronal mouse slice showing the final electrode bundle locations after up to five electrode advancements of ∼60 µm (white dots). Brain slice image credit to the Allen Mouse Brain Atlas. ***C,*** Electrode location did not significantly affect task performance (n.s. = not significant). Each circle represents performance during one session.

### MOs is more active and has more sample port-selective neurons during the sample/encoding phase of WM

We tracked subregional neural firing activity throughout all key phases of the DNMTP task. Each pseudopopulation contained neurons exhibiting task-related changes in firing rate (FR) that consistently appeared within a brief time window around the sample phase nose poke across many trials (example neuron in Fig. 2*A*). Heat maps of the FR were created by first Z-scoring the spike counts in 100 ms bins for all neurons in separate pseudopopulations, followed by sorting them from highest to lowest peak Z-scored activity around the sample nose poke (Fig. 2*B*). Quantification of mean Z-scored pseudopopulation FR at each time point revealed that the MOs significantly increased its firing rate compared to the other two implant locations in a 200-300 millisecond window before the sample poke (Fig. 2*C*, for details on statistics see **Methods**). There were no significant differences between dmPFC and vmPFC at any time points. We also quantified the percentage of neurons in each pseudopopulation that either increased or decreased their firing rate by 0.5 Z-units around the sample (Fig. 2*D*). In correspondence with the above findings, the MOs had the largest proportions of both increasing and decreasing units compared to the other regions around the sample poke, while vmPFC contained the lowest. In both figure panels, and throughout the rest of the paper, time bins with significant pairwise differences (p < .05, see **Methods**) between two mPFC subregions can be visualized in the figures as straight lines containing their two respective colors.

**Figure 2.**
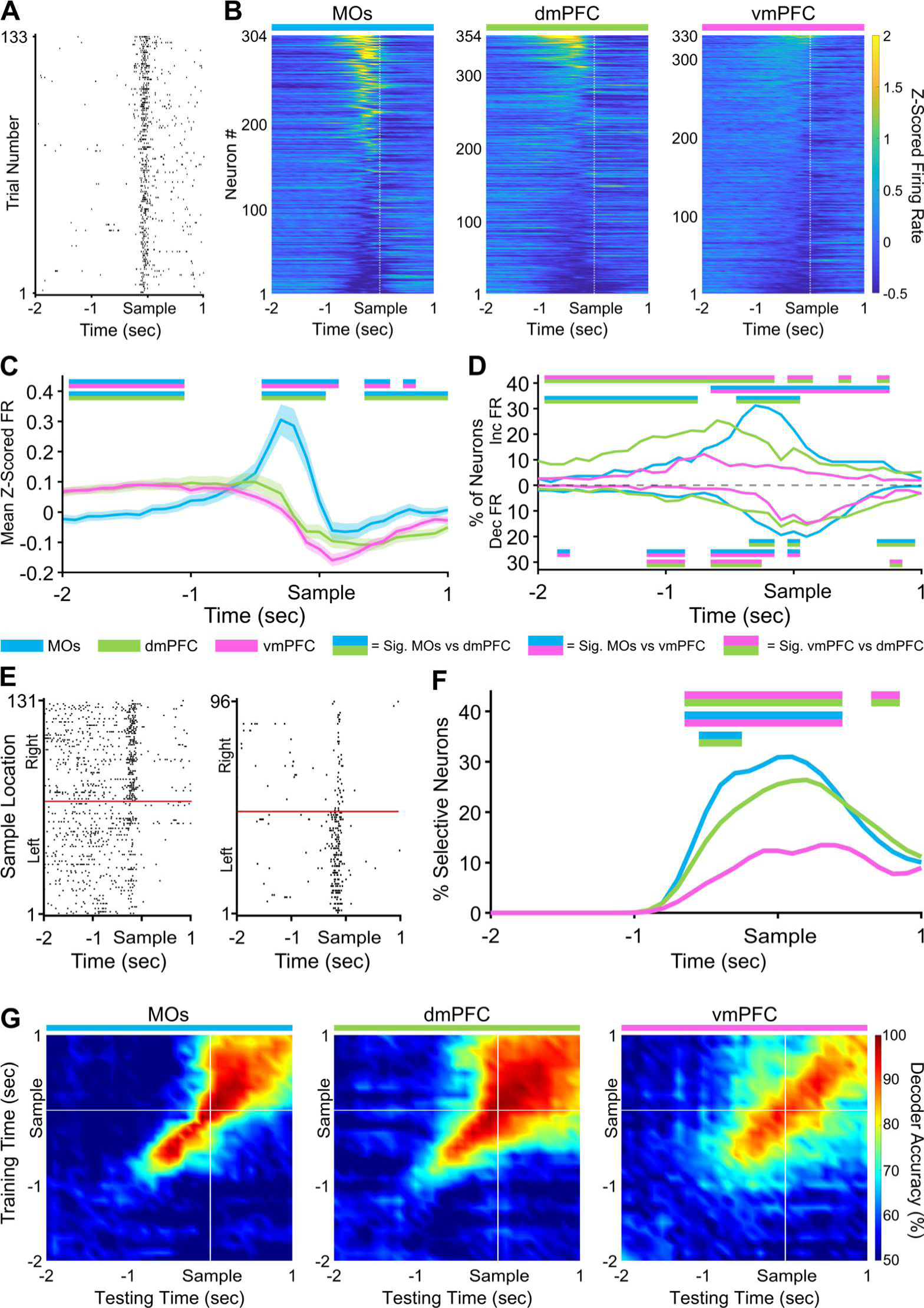
Sample location selectivity is highest in the MOs during the sample phase of the DMNTP task. ***A,*** Example raster plot from an MOs neuron increasing its firing rate around the sample port poke on all correct trials. ***B,*** Z-scored heat maps of all neurons recorded from each region around the sample poke (white dashed line) sorted by each neuron’s mean Z-scored firing rate 500 ms before the sample poke. ***C,*** The mean Z-scored pseudopopulation firing rate is higher in MOs leading up to the sample poke compared to the other regions. ***D,*** The percent of neurons either increasing or decreasing their firing rate is also higher in the MOs. ***E,*** Example sample location-sorted raster plots of two MOs neurons exhibiting transient selective firing for the right or left sample location, respectively. Red line separates left from right sample trials. ***F,*** MOs contains the highest percentage of neurons selective for a specific sample location in a ∼1 s window around the sample poke. ***G,*** Sample location decoding accuracy of a linear support vector machine trained and tested on every combination of 100 ms time bins from the sample window. White lines indicate sample port poke. In panels *C*, *D*, and *F*, double-colored straight lines represent statistically significant differences (p-value < .05) between the two respective subregions in that time bin, after correcting for both false discovery rate and family-wise error rate.

We next looked at how selective the pseudopopulations were for the sample port location in the same period around the sample poke (examples of selective MOs neurons in Fig. 2*E*). Using a permutation testing method (see **Methods** for details), we found that the MOs also contained the most neurons that could significantly differentiate sample port location based on their FR (Fig. 2*F*). This was followed by dmPFC and lastly by the vmPFC. These results suggest a functional gradient, which is strongest in MOs and weakest in vmPFC, in the extent to which these different subregions encode not only the beginning of the sample phase but also the sample port location. The stability of sample location selectivity was subsequently measured using cross-temporal linear support vector machine (SVM) decoding analysis (Fig. 2*G*, see **Methods** for details). In the MOs, the predictive ability of the decoder was nearly 100% effective within 100-200 ms of the training time bin, but this effectiveness rapidly decreased to chance levels in a few hundred milliseconds. In contrast, models trained on timepoints 0.5s before the sample poke were able to predict sample location for up to one second after the sample poke with ≥70% accuracy in both the dmPFC and vmPFC, but not in the MOs, indicating a more temporally stable population code in the former two regions.

### Retrospective sample information is stably maintained in the dmPFC throughout the entire five second delay

The approaches taken above were next applied to the delay phase of the task. MOs Z-scored population activity around the delay poke was elevated above the other subregions to a degree comparable to the sample phase (Fig. 3*C*). This poke-related difference in pseudopopulation FR between subregions did not persist into the five second delay holding period. Similar to its activity in the sample phase, the vmPFC contained fewer neurons appreciably changing their activity throughout the delay (Fig. 3*D*). Combined with the observation that the vmPFC also had a notably small number of selective neurons (Fig. 3*F*), we conclude that this subregion plays a minimal role during the delay period in our task.

**Figure 3.**
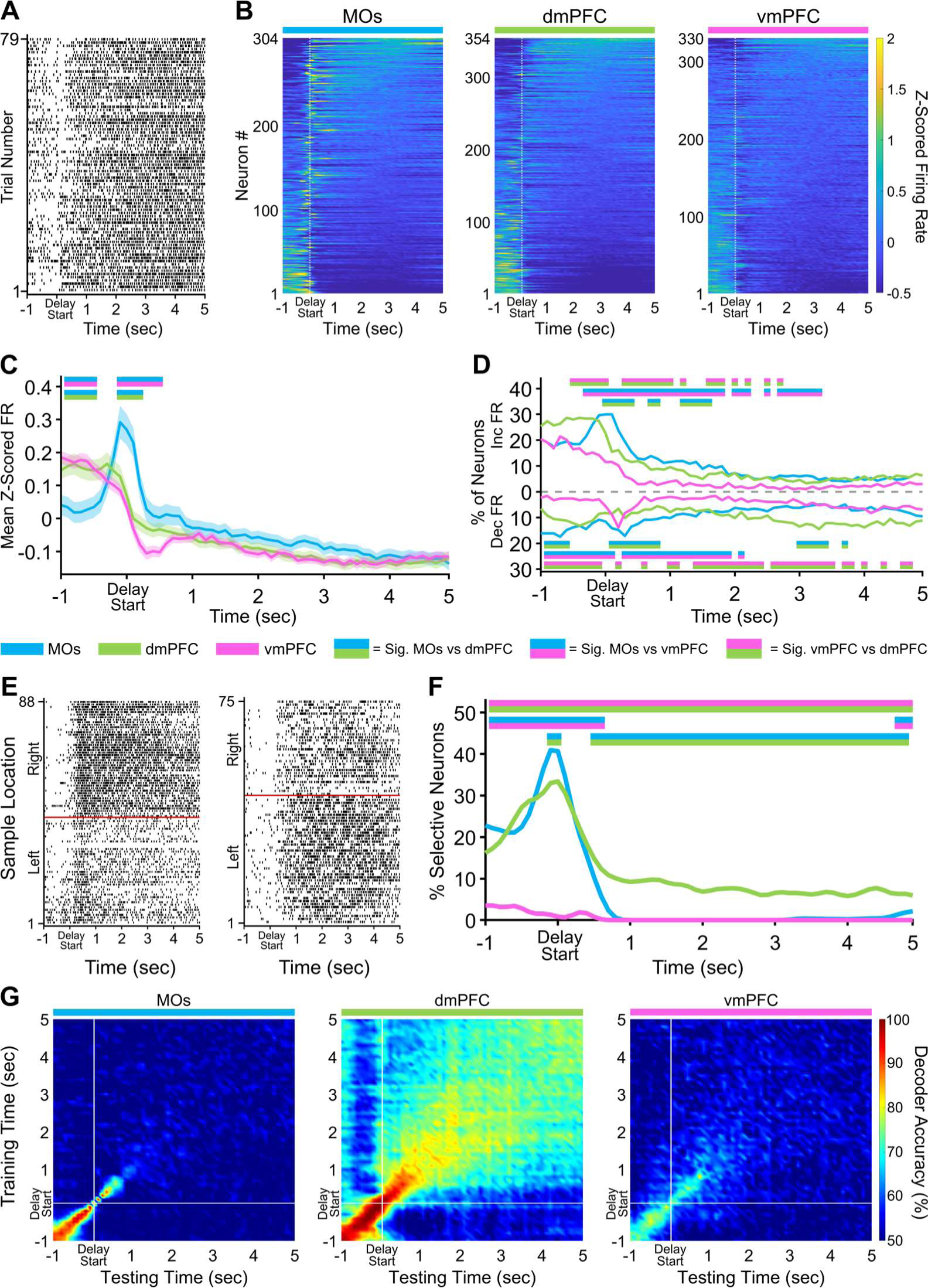
Delay activity in dorsal mPFC is persistently selective for retrospective sample location. ***A,*** Example raster plot from a dmPFC neuron aligned to the start of the delay hold period for all correct trials. ***B,*** Z-scored heat maps of all neurons recorded from each region, sorted by each neuron’s mean Z-scored firing rate during the delay phase (5 s period after the white dashed line). ***C,*** The mean Z-scored pseudopopulation firing rate is higher in MOs around the delay poke compared to the other regions. ***D,*** The percentage of neurons either increasing or decreasing their firing rate is also higher in the MOs at delay start. ***E,*** Example sample location-sorted raster plots of two dmPFC neurons exhibiting persistent selective firing for retrospective right or left sample location throughout the delay phase. Red line separates left from right sample trials. ***F,*** MOs and dmPFC contain similar percentages of neurons selective for a specific sample location in a ∼1 s window around the delay poke, but only the dmPFC shows persistent sample location selectivity throughout the entire delay phase. ***G,*** Sample location decoding using a linear support vector machine that was trained and tested on every combination of 100 ms time bins during the delay phase confirms the existence of persistent selectivity only in the dmPFC. White lines indicate the start of the delay period (back port poke). In panels *C*, *D*, and *F*, double-colored straight lines represent statistically significant differences (p-value < .05) between the two respective subregions in that time bin, after correcting for both false discovery rate and family-wise error rate.

The most striking finding during the delay phase was a time-dependent transition in the subregion that encoded the retrospective sample location most prominently. For the 200 ms around when the mice poked the back delay port, the MOs had the highest proportion of selective neurons by a small but significant margin over the dmPFC. Slightly less than one second into the delay hold, the dmPFC became the only subregion to contain *any* neurons with retrospective sample port selectivity throughout the rest of the holding period (Fig. 3*E,F*). This pattern of sample selectivity throughout the delay holding period suggests that there may be a subset of dmPFC neurons that maintain a persistently higher firing rate on either left or right sample trials. However, alternative WM mechanisms have been theorized which rely on more dynamic representations involving chains of multiple different neurons becoming transiently selective at different points in time, leading to an unstable population code^38^. Using the same cross-temporal SVM analysis as before, we were able to infer that the pattern of activity was likely due to a persistent mechanism. Decoding accuracy remained above 70% from the beginning to the end of the delay holding period in the dmPFC suggesting that a stable population code was present during this time (Fig. 3*G*). Conversely, the MOs and vmPFC decoded at chance levels throughout most of the delay.

### Retrospective sample port information is most strongly encoded by the dmPFC before and after the choice poke

Reminiscent of nose pokes in the previous phases, the MOs was again the most active subregion directly around the choice poke (Fig. 4*C,D*). We also recorded a large amount of activity starting ∼1.5 seconds before the choice poke which appeared to be highest in the vmPFC, although this difference was not statistically significant in this time frame (Fig. 4*C*). Contrary to the previous two task phases, the selectivity in this pre-poke period and the period directly around the poke was most represented in the dmPFC instead of the MOs (Fig. 4*E,F*). This significance continued in dmPFC until about one second after the choice, once again implying that the dmPFC stably represents the location of the sample port that was visited earlier in the trial. Consistent with this idea, the cross-temporal SVM uncovered a more stable code both before and after the choice poke in dmPFC compared to other areas (Fig. 4*G*).

**Figure 4.**
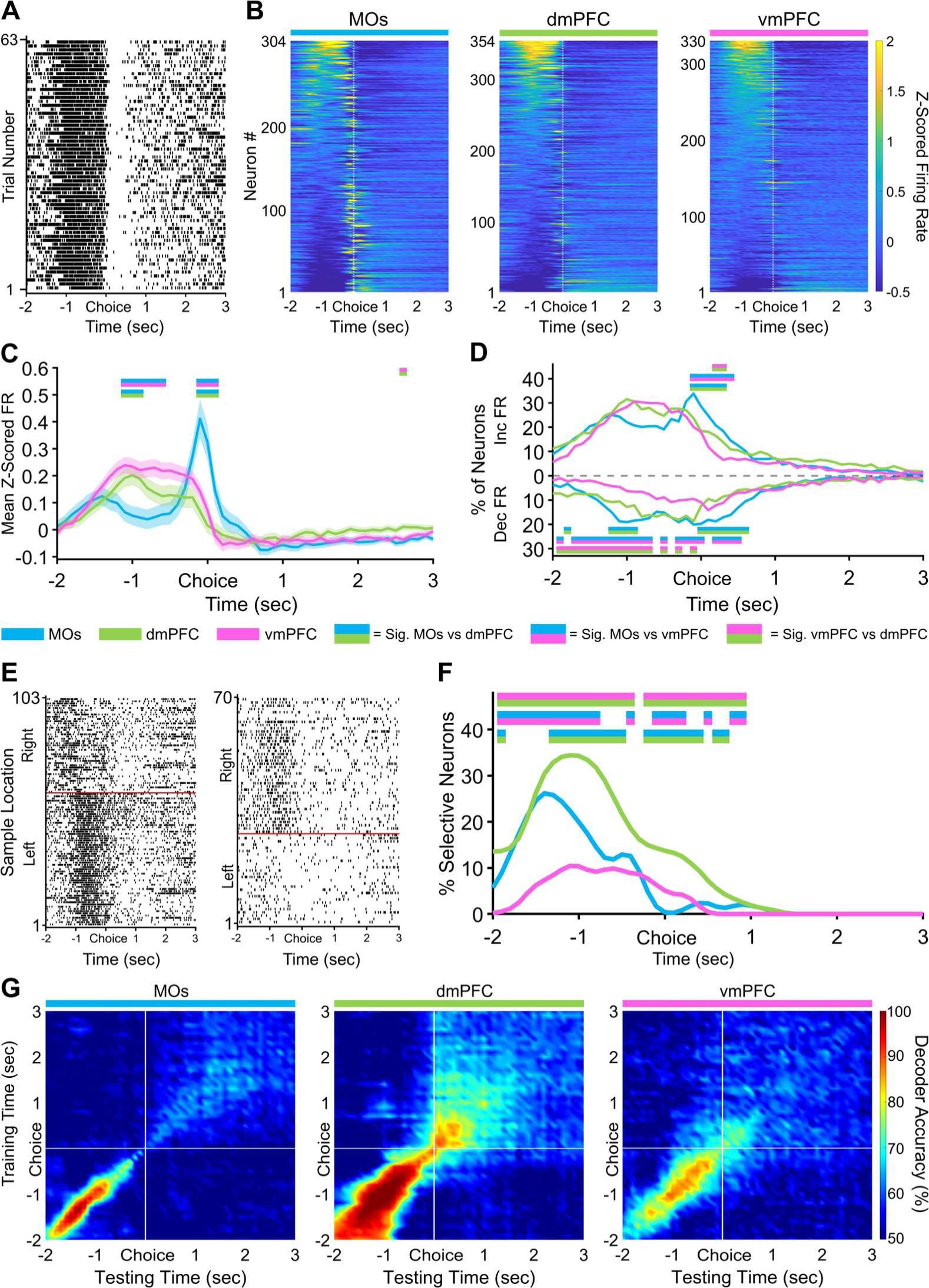
During the choice phase, vmPFC has a higher pseudopopulation pre-choice Z-scored firing rate, and dmPFC contains more neurons selective for retrospective sample location. ***A,*** Example raster plot from a vmPFC neuron aligned to the non-match choice poke for all correct trials. ***B,*** Z-scored heat maps of all neurons recorded from each region, sorted by each neuron’s mean Z-scored firing one second before the choice poke (white dashed line). ***C,*** The mean Z-scored pseudopopulation firing rate is higher in vmPFC leading up to the choice compared to the other regions, while activity right around the choice poke is highest in MOs. ***D,*** The percentage of neurons increasing their firing is higher and the percentage of neurons decreasing their firing rate is lower before the choice in vmPFC. ***E,*** Example sample location-sorted raster plots of two dmPFC neurons exhibiting transient selective firing for the right or left sample location during the delay phase. Red line separates left from right sample trials. ***F,*** Although activity is higher in vmPFC leading up to the choice poke, dmPFC contains the most retrospective sample port selective neurons out of all three regions. ***G,*** Sample location decoding accuracy of a linear support vector machine trained and tested on every combination of 100 ms time bins during the non-match choice phase. White lines indicate when the mice poked the choice port. In panels *C*, *D*, and *F*, double-colored straight lines represent statistically significant differences (p-value < .05) between the two respective subregions in that time bin, after correcting for both false discovery rate and family-wise error rate.

### MOs most strongly differentiates task phase

Although selective coding of retrospective sample port identity is necessary for successful DNMTP performance, it is not the only task parameter that mice must constantly monitor to make the correct non-match choice and maximize reward. A mouse must also be aware of several other aspects of the task including tracking which phase of the task it is in so that the correct motor strategy can be enacted at specific points in time, upcoming choice port location and the location of the choice it eventually makes, and what the outcome of the choice was (correct or incorrect) so that the mouse can update its strategy on the next trial if necessary. Any or all of these factors may interact at any given point throughout the time course of a single trial, so we wanted to analyze each factor with respect to the other factors.

For this, we used a general linear model (GLM) to predict how much the firing rate of each neuron in 200 ms bins around pokes depended on the four following predictor variables: poke context (sample, delay, or choice poke), sample port location (left or right), choice port location (left, center, or right), and outcome (correct or incorrect). Shuffled coefficient of partial determination analysis (CPD, see **Methods** for details) unveiled the percent of total firing rate variability that was explained by each of these variables. For this first GLM analysis, we were mainly interested in how closely the PFC monitored task phase information around pokes. The task phase the mouse was transitioning into around a poke accounted for significantly more total neural variability in the MOs as compared to dmPFC and vmPFC (Fig. 5*A*). The dmPFC furthermore encoded this variable to a larger degree than vmPFC. Poke context accounted for an overwhelming percentage of all *explainable* variability in each region (Fig. 5*B*), confirming that information about task progression is a crucial part of PFC computation regardless of the subregion. In lieu of these results, we assessed how well the population activity in separate subregions could decode the poke context around poke events. We implemented a similar SVM approach as the one described in previous sections, but we applied a multiclass coding scheme instead of a binary one since the model now had three possible choices (sample, delay, or choice) to differentiate. This method was able to decode poke context with 100% accuracy in all subregions (Fig. 5*C*).

**Figure 5.**
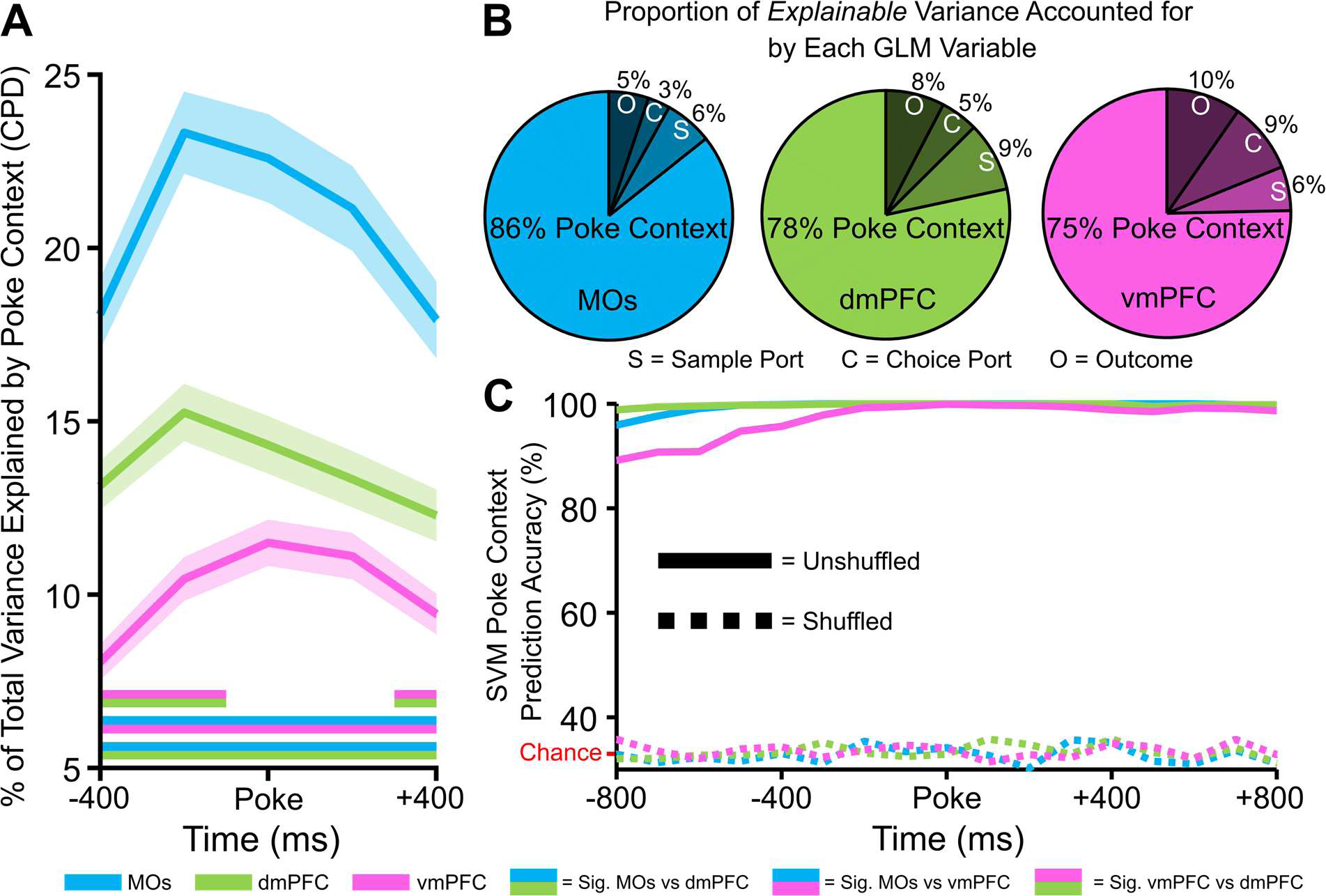
Encoding of poke context (sample, delay, or choice poke) accounts for the large majority of explainable firing rate variability in all three regions. ***A,*** The coefficient of partial determination (CPD) from a general linear model was used to find the fraction of total firing rate variance around pokes that can be explained by the poke context. The CPD is highest in the MOs pseudopopulation. Double-colored straight lines represent statistically significant differences (p-value < .05) between the two respective subregions in that time bin, after correcting for both false discovery rate and family-wise error rate. ***B,*** CPD was also calculated for the sample port, choice port and outcome (correct vs incorrect trials) variables. When combined with these other three predictor variables, poke context variability accounts for over 75% of explainable firing rate variance in all regions. ***C,*** Poke context is decodable with nearly 100% accuracy in all regions using a linear support vector machine (SVM). Chance level decoding in the shuffled control (dashed lines) is 33%.

### GLM exposes distinct subregional processing of different task variables

In order to investigate the contribution of DNMTP task variables over the same wider time frame used in the first several figures, we needed to remove the poke context task variable due to its dependence on a prohibitively small window around pokes. Therefore, three predictor variables remained for this GLM over time: sample port location, choice port location, and outcome. With a similar CPD approach as above, we calculated the percent of neurons in each region that significantly (according to a shuffled CPD control analysis, see **Methods** for details) encoded the specified task variables in each 200 ms bin. This produced analogous results to our retrospective sample location selectivity investigation using permutation testing in Figs. 2, 3, and 4, further strengthening these findings (Fig. 6*A*). The time-based GLM also established that neither upcoming choice port identity nor trial outcome were encoded in the sample or delay phases of the task (Fig. 6*B,C*). Importantly, we only observed a considerable number of choice-port-selective neurons around the choice poke itself, with the vmPFC displaying the largest percentage (Fig. 6*B*). Likewise, only after the choice was made were we able to find significant subregional differences in the number of neurons encoding outcome (Fig. 6*C*). This was a compelling affirmation that the mice were not choosing an incorrect prospective motor plan or specific choice location before they needed to make the actual choice. Surprisingly, the MOs was the region with the most neurons encoding the outcome variable, possibly due to motor activity changing drastically on correct vs incorrect trials as mice were either consuming a water reward by licking (correct), or removing their nose from the port when no water was dispensed (incorrect).

**Figure 6.**
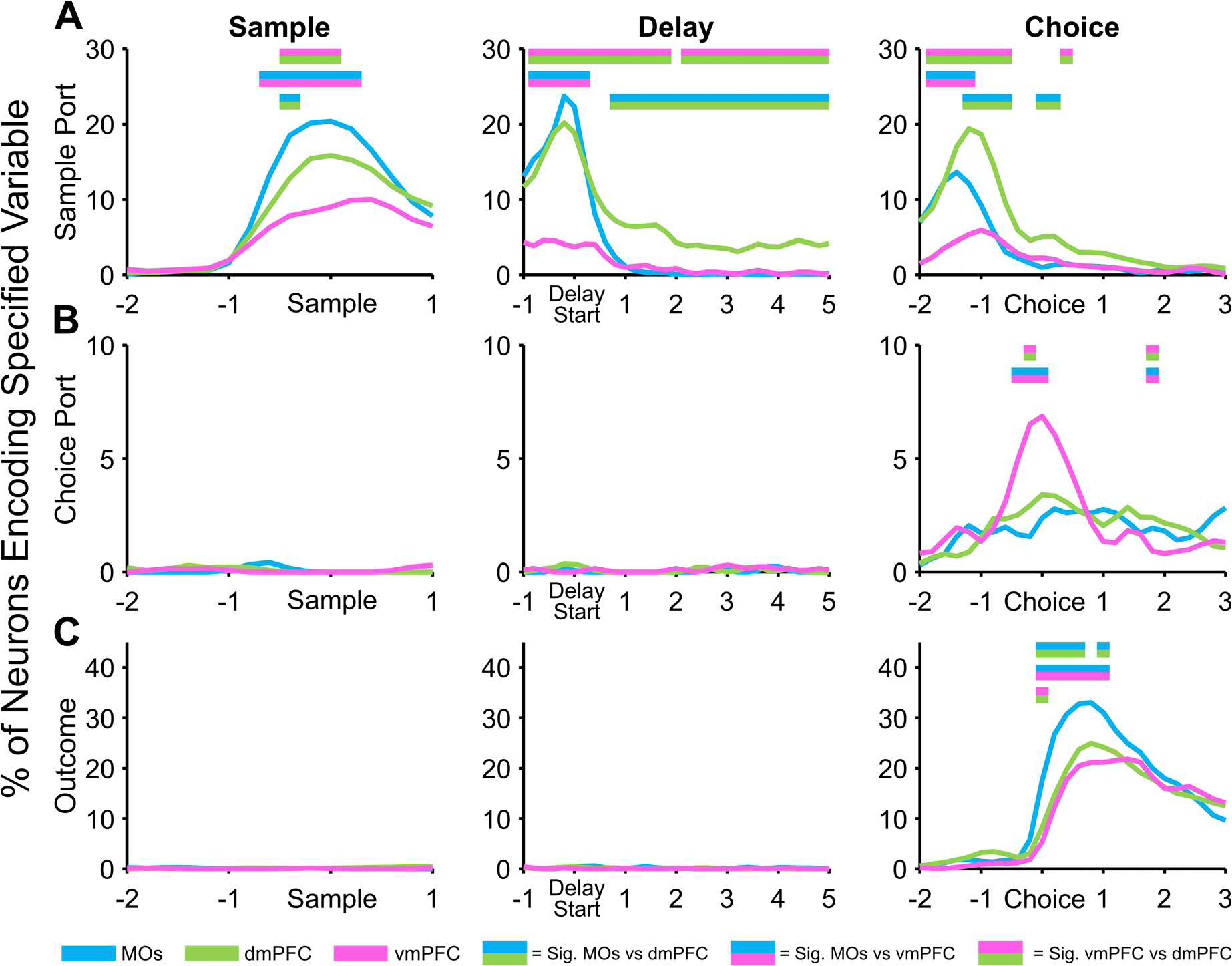
General linear model (GLM) reveals PFC subregion-specific patterns of information encoding across task phases of a working memory task. After removing the poke context regressor, a GLM uncovers the percent of neurons in each region that significantly encode working memory task variables (sample ports, choice ports, and outcome (correct vs incorrect)) at key timepoints. ***A,*** The region with the highest percentage of significant sample location neurons shifts from MOs to dmPFC as the mice progress through task phases. ***B,*** Prospective choice port location is not detectible in any region until the choice phase, during which vmPFC shows the highest selectivity. ***C,*** The trial outcome (correct vs incorrect), is similarly not encoded by any region until the choice is made. Each region encoded the outcome (correct vs incorrect) after the choice was made, with the MOs having the largest proportion of outcome-encoding neurons. Double-colored straight lines in any panel represent statistically significant differences (p-value < .05) between the two respective subregions in that time bin, after correcting for both false discovery rate and family-wise error rate.

### GLM beta weights for retrospective sample location selectivity confirm its representational stability throughout trials

Since sample location was the only task feature significantly encoded by all subregions throughout all phases of WM, we wanted to take a closer look at the temporal dynamics of this information in each area. We reasoned that knowledge about the way in which retrospective representations evolve is critical to fully understand the circuit mechanisms underlying WM, as most definitions of WM rely on the ease with which animals can maintain previously encoded information over time. A recurring theme in our data is that this sample location information seems to be strongly reactivated very close to poke events. We therefore investigated the extent to which the sample selectivity of neurons in each region was reactivated around these pokes. The percentages of neural selectivity that was significant across multiple pokes was quantified in the Venn diagrams in Fig. 7*A*. MOs and dmPFC both had a large number of neurons selective for the sample location across multiple task phases. Interestingly, there were few vmPFC neurons that were sample-selective over multiple pokes.

**Figure 7.**
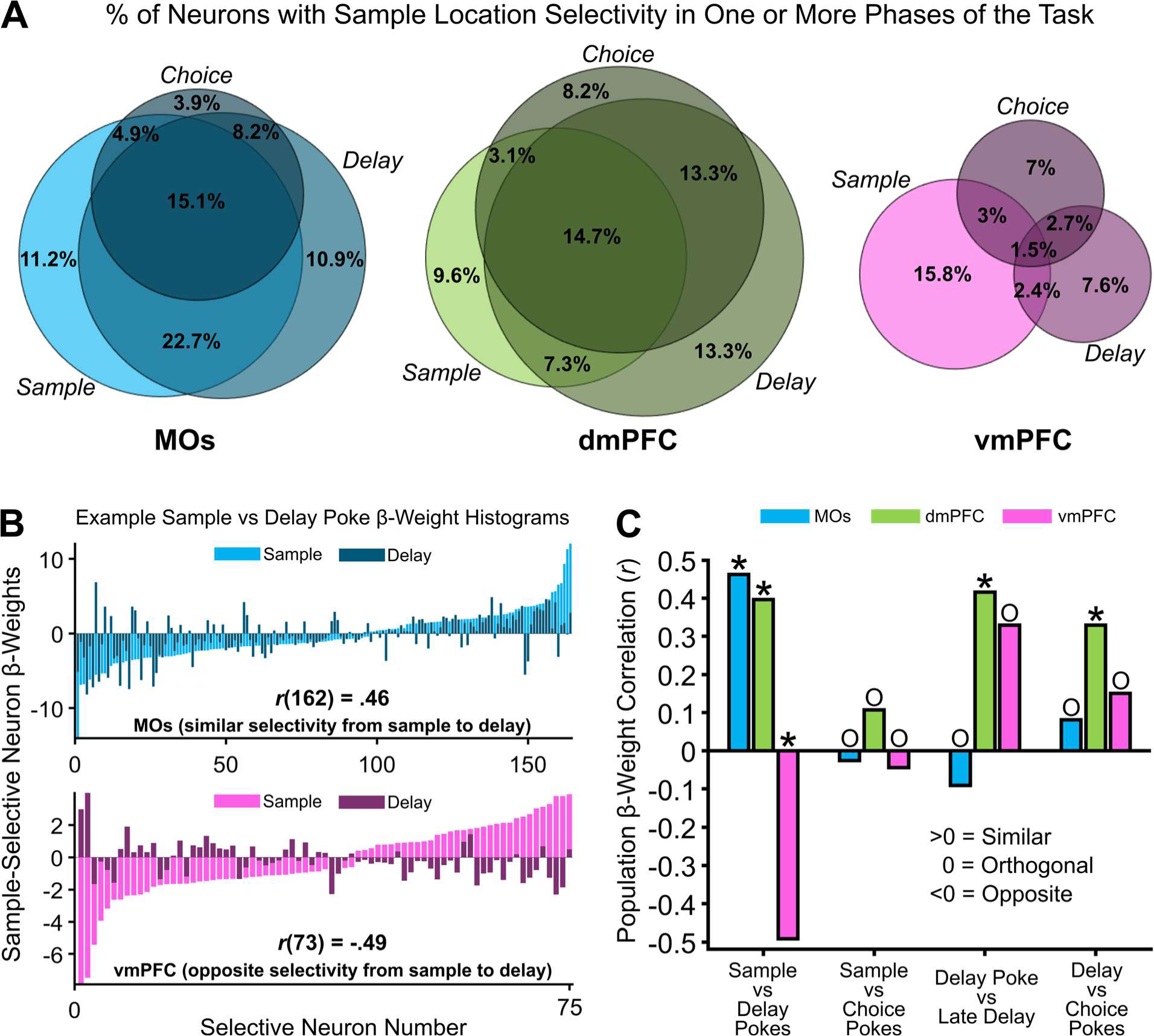
Sample location selective firing rate representations have higher stability in MOs and dmPFC compared to vmPFC as the mouse progresses through working memory task phases. ***A,*** Venn diagrams depicting the percent of neurons in each region that were selective for the sample location around the poke initiating each task phase. Overlapping circles in the Venn diagram represent the percent of neurons that share retrospective sample location selectivity around either two or all three task phase pokes. ***B,*** Example GLM-derived beta weight histograms from significant sample location-encoding neurons in the GLM showing how most neurons in the MOs remain selective for the same port across sample and delay pokes (top), but neurons in the vmPFC switch initial sample location selectivity from sample to delay (bottom). ***C,*** Pearson beta weight correlations quantifying the similarity in location selectivity between pokes initiating one task phase, to pokes initiating another one. MOs location selectivity is similar from sample to delay pokes, but destabilizes over the course of the task. dmPFC remains the most stable across time. vmPFC shifts location selectivity from the sample to delay pokes, but then stabilizes later in the task. Asterisks represent an *r* value that is significantly different from zero, while O (<u>O</u>rthogonal) represents an *r* value that is not different from zero. The reason for the non-significant *r* value from the vmPFC correlation in the *Delay Poke vs Late Delay* correlation is because of the much smaller number of selective neurons in vmPFC during this time.

The maximum GLM beta weights from these selective neurons were identified around pokes (600 ms before and after poke). We sorted these beta weights by amplitude for better visualization, and plotted examples of these histograms for MOs and vmPFC in Fig. 7*B*. In these examples, negative beta weights represented a neuron with left sample selectivity, while positive signified right selectivity, and higher amplitude meant the neuron had a higher average difference in firing rate between left and right trials. To check temporal stability of these beta weights, we ran simple Pearson correlations between beta weights from significantly selective neurons at two different time points – for example, sample poke (light bars) versus delay poke (dark bars) as shown in Fig. 7*B*. The *r* values from these correlations are graphed as bars in Fig. 7*C* for every subregion across every time-based comparison. Briefly, we found that the sample-selective subpopulations of MOs and dmPFC neurons have similar beta weight distributions when comparing sample and delay pokes, while the vmPFC subpopulation is negatively correlated. The dmPFC also had similar selectivity across all adjacent time comparisons, adding more evidence that it is uses the most stable coding of all three subregions. Interestingly, none of the areas were stable from sample to choice pokes, which potentially arises from the fact that these events are separated in time by a cognitively demanding delay phase, which may largely reorganize neural activity.

### vmPFC Z-scored population activity displays the largest change in reward outcome-related firing rate

Up to this point, we had not detected any substantial contributions from the vmPFC to DNMTP WM performance. However, this region is known to have the densest reciprocal connections with the ventral tegmental area and amygdala out of all PFC subareas^39,40^, hinting at potential involvement in more valence or reward-based information processing. As a result, we subtracted the mean Z-scored firing rate of all recorded neurons on correct trials from that on incorrect trials (subsampling for the lower count of incorrect trials), which gave rise to more pronounced differences in pre-choice and post-outcome population activity in the vmPFC compared to MOs and dmPFC (Fig. 8*A*, positive values indicate higher FR on incorrect trials). Not surprisingly, the vmPFC was also different relative to MOs and dmPFC in terms of the percentages of neurons responding by either increasing or decreasing their firing rate based on the reward outcome (Fig. 8*B*).

**Figure 8.**
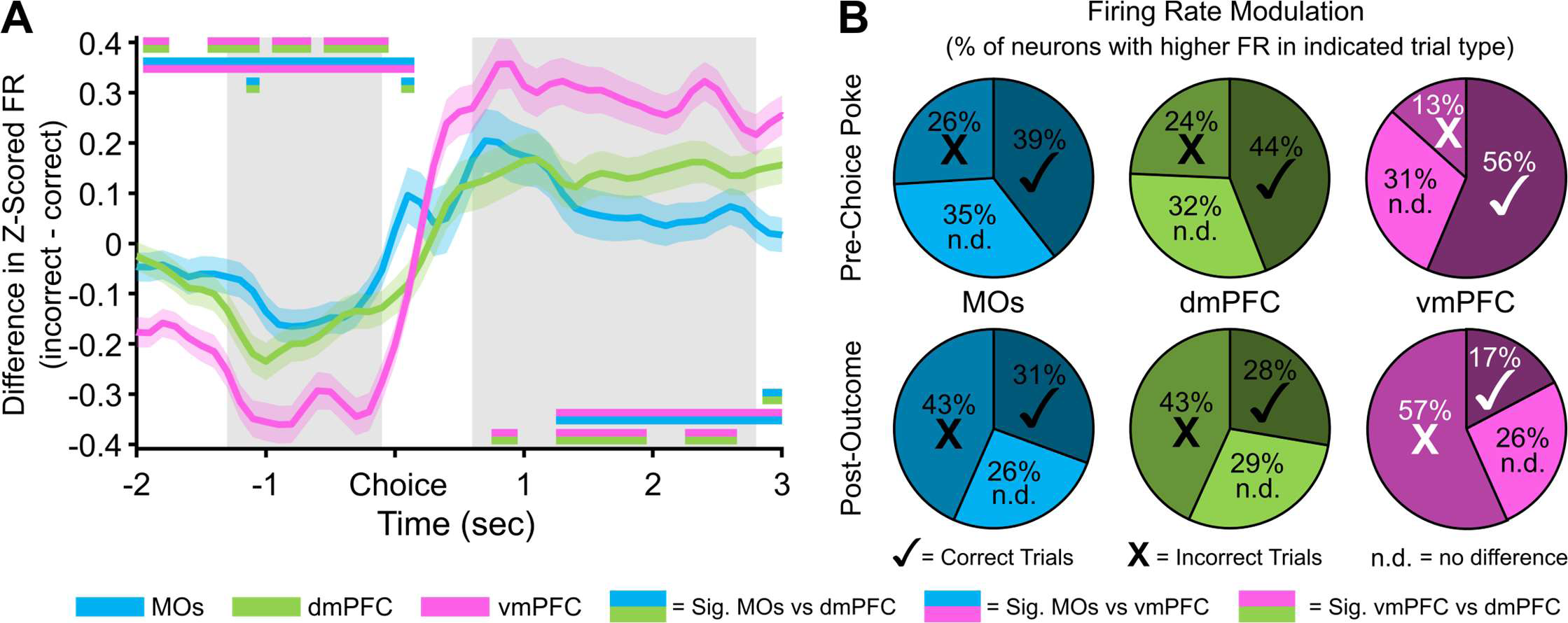
Differences in pseudopopulation activity between correct and incorrect trials are more pronounced in the ventral mPFC before and after the choice poke. ***A,*** Z-scored firing rate differences between correct and incorrect trials in each region over time. vmPFC shows the largest decrease in activity on incorrect trials before the poke, and the largest increase in activity in response to an incorrect outcome after the choice. Grey bars represent the time windows used for the pre-choice and post-outcome calculations in panel B. Double-colored straight lines represent statistically significant differences (p-value < .05) between the two respective subregions in that time bin, after correcting for both false discovery rate and family-wise error rate. ***B,*** Proportion of neurons in each region that have a higher Z-scored firing rate on Correct (Π) or Incorrect (**X**) trials, or showed no difference in Z-scored firing rate between correct and incorrect trials (n.d.). The proportion of neurons modulating their firing rate on correct vs incorrect trials is also greater in vmPFC. White marks instead of black represent statistically significant proportions of the neurons with higher firing rate on that trial type in the vmPFC compared to the other 2 regions. There were no differences between the other regions.

## Discussion

In this paper, we delineate how the neural populations in adjacent mouse mPFC subregions processed information about WM task-related variables over time. The first key finding was that each subregion exhibited characteristic and differentiated activity during the task relative to the other subregions. The second key finding was that much of the WM task-related activity appeared unrelated to storing WM information (i.e., representations of the previously visited sample port locations). Rather, it reflected changes in activity related to poke context (sample, delay, or choice poke), and rewards. The third key finding was that WM of the sample location was stored briefly (∼1 second) in the MOs and dmPFC and in a sustained manner only in the dmPFC, with minimal retrospective representations detected in vmPFC. The final key finding was that vmPFC predominantly represented choice and outcome-related activity relative to MOs and dmPFC.

Of note, there is a dearth of comparative analysis using single neuron and population activity of neighboring mPFC subregions during WM, particularly in rodents. Activity across mPFC subregions has been examined in rats performing DNMTP tasks^41,42^ and a more complex match to place task^43^. In these studies, single neurons responded prominently to reward locations and task phase, but in contrast to our findings in this study, no sustained neural activity related to retrospective sample port location was noted during the delay period. Expanding on these studies, our work implements complementary computational analyses to further explore the disparate functional roles and temporal dynamics/stability of subregional mPFC neural populations during WM. In addition, we included another brain region in our experiments, the MOs, after recent evidence that it supervises more abstract functions than basic motor control and should be considered a part of the mPFC^44^.

In our DNMTP task, the MOs behaves in a manner consistent with the theory proposed by Barthas and Kwan (2017) which posits its involvement in context-dependent selection of motor plans^45,46^ and the online monitoring of sequential sensorimotor tasks^47^. Activity in our recorded MOs neurons peaks in a tight window around all port pokes. Importantly, population-level differences in this transient activity can simultaneously differentiate which part of the task the poke is occurring in alongside distinguishing the left versus right sample port location. These transient poke-centered phenomena may represent close monitoring of the phase of the DNMTP task the animal is in at fixed intervals, while also providing information at each stage about the spatial rule of the current trial, facilitating ongoing motor plan selection and timing. Consistent with the notion that MOs is involved in context-dependent selection of motor plans, it has been reported that MOs can relay its contextual/motor information to many areas of the neocortex it connects with, including sensory and primary motor cortex^48^. Future timed inactivation studies may be useful in exploring these ideas further in WM tasks.

With respect to retrospective sample location selectivity, our analyses uncovered strong sample port representations in both MOs and dmPFC during all task periods. The retrospective sample location was detectable using single neuron and population analyses, including using an SVM to decode the sample port location and developing a GLM to show that this port location information is significantly encoded in MOs and dmPFC neurons well after the sample phase. As mentioned above, the retrospective sample location-related activity in MOs manifests only briefly around pokes (∼1 sec), and these recurring transient patterns of selectivity do not remain stable throughout a trial. We speculate that MOs may not contain the molecular or circuit architecture necessary to maintain persistent activity in a group of neurons^49^. In contrast, subsets of dmPFC neurons are selective for the retrospective sample port at the sample, delay and choice pokes in similar patterns, with a smaller group (∼10%) exhibiting sustained retrospective sample port selectivity throughout the delay. These findings are consistent with the notion that there are overlapping ensembles of WM neurons operating with different dynamics in different PFC subregions on different timescales. They also suggest that briefly active WM ensembles may be more common than sustained ensembles. It will be useful to explore these ideas further in primates and rodents using different WM tasks with a range of delay lengths.

Perhaps the most intriguing finding in our data is the aforementioned stability of dmPFC retrospective sample port selectivity of a group of neurons throughout the course of behavioral trials, especially during the majority of the delay holding period when neither of the other subregions display sample location-related activity. A substantial body of evidence has set the precedent for the existence of persistently selective delay representations in various primate brain areas including dorsolateral prefrontal cortex^50,51^, and in head-fixed mice. The latter work was done in the nearby anterior lateral motor cortex (ALM) during a delayed motor response task showing sample-selective preparatory activity for an upcoming left or right lick^30–32,52,53^. This task differs from ours in that the head-fixed animals know the exact location of the correct choice throughout the delay period, allowing them to make a precise motor plan which may be represented as persistent activity. In our task, the prospective location that will be rewarded cannot be anticipated. Additionally, the delay lengths used in the reports cited above are relatively short (1-2 seconds) compared to ours at five seconds. We believe that this longer interval is crucial to unraveling the dynamics of WM, as we observed that most neurons coding for the retrospective sample location in MOs and dmPFC around the delay poke are not sustained beyond 1-2 sec into the delay, with a small group of neurons (∼10%) in dmPFC exhibiting sustained WM activity. Other studies in freely-moving rats also found non-sustained sample-selective delay activity in the frontal orienting field during a WM motor planning task^45^, and in prelimbic cortex in a delayed alternation task^54^.

An often-overlooked aspect of WM is the need to maintain some signal which updates strategies based on feedback from reward outcomes, so that behavior can be adjusted in the immediate future if results don’t match expectations. Our findings in vmPFC align with the possibility that this region is predominantly involved processing choices and outcomes. This could be communicated by different release patterns of dopamine in this region ^55^, or changes in firing from glutamatergic amygdala inputs ^56^. Overall, our results point toward a dynamic flow of information from MOs to vmPFC as the mice progress through the task. Similar dynamics across distant brain regions have been characterized before in humans ^22^, monkeys ^57^, and mice ^30^, but they remain poorly characterized both in primates and mice in PFC subregions involved in WM.

One caveat of this study is that we did not record from these subregions simultaneously, due to the technical difficulty of probing multiple sites along the curvature of the cortex. Future studies would benefit from using more advanced electrophysiological setups (e.g., Neuropixels 2.0 probes^58^) to record all subregions at once. We also did not collect video data to accompany these recordings. In future studies, video would be useful for connecting specific movements to associated neural activity patterns^59^. Furthermore, with the way our task was designed, we cannot definitively determine if the mice are actively remembering the retrospective port location or prospectively planning a left or right turn *away* from the sample port during the delay period^60^, although both of these scenarios require a WM mechanism. However, our GLM data showing no neurons in any region that were selective for the upcoming choice port lends support to the idea that the sustained representation we describe is most likely a retrospective one that aligns well with stored WM information very closely related to the sample port visited earlier in the trial.

One ongoing challenge in neuroscience is the mapping of rodent mPFC subregions to the corresponding areas from primate PFC. Although this can be done according to common afferent and efferent projections, molecular expression, and functions^61–64^, there is often a lack of consensus regarding the extent of similarity/homology. Another challenge has been the difficulty in eliciting persistent WM activity in rodent PFC to be able to study the underlying mechanisms in mice. By identifying such activity, as well other types of WM activity, this study provides a foundation to perform such studies using tools that are uniquely available in mice. Our work here also paves the way for more detailed analysis of these networks and a more nuanced and dynamic view of how neighboring brain regions process information in complex and complementary ways as mice progress through a complicated behavior that requires WM and other functions.

## Methods

### Animals

All animal experiments performed in this study were approved by the Veterans Affairs Portland Health Care System Institutional Animal Care and Use Committee. Nine female and seven male mice, bred on a C57BL/6J background, were housed in the Veterans Affairs Portland Health Care System Veterinary Medical Unit on a reverse 12-hour light cycle with lights turning off at 8:00 AM (PST), and on at 8:00 PM (PST). All mice were group-housed before electrode implantation, after which they were single-housed until the completion of the experiment to prevent them from damaging each other’s implants. Mice had *ad libitum* access to water (unless restricted for experimentation) and PMI PicoLab 5L0D Laboratory Rodent Diet (LabDiet, Inc., St. Louis, MO, USA). Mice were housed in rooms with constant temperature (22-26 °C dry bulb) and humidity (30-70%) monitoring.

One day prior to the start of initial behavioral training, mice were water restricted to 85-90% of their initial body weights so that they were motivated to seek out water rewards. This weight-based water restriction continued for the duration of active behavioral experimentation (but not during recovery from surgery), after which they were promptly returned to *ad libitum* water until they were sacrificed for implant location confirmation. The long-term water restriction protocol consisted of giving the mice about one gram of a 98%-water, gelatinous hydrocolloid mixture (HydroGel^®^, ClearH20, Westbrook, ME, USA) daily, after behavioral tasks, to keep the mice at a constant water-motivated weight. Furthermore, during the one-week recovery from surgery, mice were supplemented with an electrolyte-based recovery diet (DietGel^®^ Recovery, ClearH20, Westbrook, ME, USA).

### Behavioral setup and training

On the first day of delayed-non-match-to-position (DNMTP) behavioral training, water-restricted mice were acclimated to the behavioral chamber (Fig. 1*A*, Bpod, Sanworks LLC, Rochester, NY, USA). The main hardware components of this chamber included a state machine control board and illuminable, photo-gated, water-dispensing ports. Integration of this system with MATLAB software (MathWorks, Natick, MA, USA) and our electrophysiological recording system allowed for precise closed-loop control over specific DNMTP task parameters via custom MATLAB scripts. These parameters included the timing of task phases, lighting of ports, delivery of water rewards or signaling of incorrect behavior with negative reinforcers, and online synchronization with electrophysiological data for accurate timestamping of neural firing and important behavioral events.

After a 15-minute habituation session on day one, mice were taught on day two that only lit ports could dispense water rewards. To do this, we randomly lit and pre-baited one of the four (three “front” and one “back”) ports so that water was available as soon as the port light turned on. This allowed mice to initially learn the simple Pavlovian association that water was available from lit, but not dark, ports. Once the mice were familiar with this association (usually after one 15-minute session), we changed to a slightly more instrumental design where water was not dispensed until the mice poked their nose in the lit port and broke the plane of the infrared photogate, prompting them to learn that their active engagement with the port was required for water to be dispensed.

The next step in training was a modified version of the final DNMTP task with intertrial intervals (ITIs), sample phase, delay phase, and choice phase. After a five second ITI, either the left or right front port had a 50% chance to randomly light up on a given trial (sample phase), and the mouse was required to poke its nose in the lit port to get a small water reward (3 µL) and activate the back delay port. The mouse then turned around to poke in the back delay port, which dispensed a one µL reward immediately after the poke. Poking in any of the dark ports during the sample or delay phases led to a punishment consisting of illumination of a bright house light and a behavioral timeout for 15 seconds, after which the exact same trial restarted. Successful progression through the delay led to the choice phase, where the previous sample port lit up along with one of the other two front ports (randomly, 50% chance). The mouse had to poke the lit port that it did not previously visit during the sample phase to complete a correct non-match choice and get a larger seven µL reward. Consumption of this reward typically took about two to three seconds, after which the mice left the port. Leaving the port after water consumption initiated a five second ITI period in which the mouse could not enter any other ports, or the ITI timer would reset. An incorrect choice similarly led to an illumination of a bright house light and a fifteen second timeout, but in this case a new trial with new port locations started afterwards.

Once mice achieved 70% non-match performance on this training task with no delay, we removed the sample phase reward and implemented a slow walk-up of delay length in each session to help the mice get to the final delay period of five seconds. The mice had to hold their nose in the back port until the delay timer ended to get the small one µL reward and enter the choice phase. After each successful delay hold, the timer went up from zero by 0.10 seconds until a five second delay was reached. We let the mice do this until they reliably got to 5 seconds for the delay period. The final version of the task had the mice starting the delay at zero seconds and walking up by one second per trial to the final five second delay. These first five trials were removed from analysis. The final task sessions, in which we also recorded brain activity, usually lasted around one hour and the mice completed anywhere from 48 to 141 trials, depending on motivation for water based on hydration status. Once mice performed at >70% for three consecutive sessions they were implanted with electrodes.

### Electrode implantations and single-unit recordings

Custom 28-channel implantable electrode bundles were constructed in-lab. The process consisted of threading 32-channel electrode interface boards (EIB-36-Narrow-PTB, Neuralynx, Inc., Bozeman, MT, USA) with 12 µm diameter tungsten wire (California Fine Wire Company, Grover Beach, CA, USA), and securing the wires in the board with gold pins (Neuralynx, Inc., Bozeman, MT, USA). Four slightly larger diameter local field potential wires were implanted in various mPFC-connected brain regions, although none of the data collected with these wires was used for analysis. Silver ground and reference wires were also soldered onto the EIB. The apparatus was built on a custom 3D-printed scaffold (Grey V4 resin, Formlabs, Somerville, MA, USA), with holes for drivable screws (McMaster Carr, Elmhurst, IL, USA) that allowed for advancement of the electrodes after every recording session. The wires were affixed to the EIBs using dental cement (UNIFAST Trad, GC America Inc., Alsip, IL, USA) to protect them from damage.

For implantation, mice were lightly anesthetized with 3% vaporized isoflurane (Covetrus, Dublin, OH, USA) and transferred to a stereotaxic surgery apparatus (David Kopf Instruments, Tujunga, CA, USA) where they were kept at ∼1% isoflurane for the remainder of the surgery. Body temperature was monitored with a Physitemp TCAT-2LV temperature controller system, which held animals between 36 and 37 °C. Prior to initial incision, they were injected with carprofen and dexamethasone for pain management, along with a topical application of lidocaine to the skull and surrounding skin. Electrode bundles were implanted on the left side of the skull at the following coordinates (from bregma): **supplementary motor area (MOs):** +1.80 mm anterior, - 1.50 mm lateral left, -1.00 mm ventral to brain surface; **dorsomedial prefrontal cortex (dmPFC):** +1.80 mm anterior, -0.40 mm lateral left, -0.50 mm ventral to brain surface; **ventromedial prefrontal cortex (vmPFC):** +1.80 mm anterior, -0.40 mm lateral left, -1.70 mm ventral to brain surface. A larger diameter reference wire was implanted in the left **striatum:** +0.50 mm anterior, -1.60 mm lateral left, -2.50 mm ventral to brain surface. A ground screw was also placed in the skull over the right cerebellum. The full setup was secured to the skull using the same dental cement mentioned previously. This included threading screws into skull-secured acrylic cuffs so that they could be advanced and drive the electrodes deeper into the brain.

Mice were allowed to recover for at least one week with daily health monitoring before returning to water restriction and behavioral testing. Electrical recordings during behavior began after two to three re-habituation sessions while plugged into the electrophysiological tether and commutator (Doric, Quebec, Canada). Data was collected with a CerePlex Direct neural acquisition system connected via a 32-channel CerePlex µ (mu) headstage (Blackrock Neurotech, Salt Lake City, UT, USA) to the implanted EIB. Unfiltered data was sampled at 30 kHz throughout an entire behavioral session. Single-units were isolated offline using Kilosort3 and timestamped to behavioral events. Since the geometry of our bundle was unknown, we used a random linear arrangement for our probe configuration parameter. We also turned off the option for registration and drift correction so that Kilosort would not try to re-order the channel map and match templates based on the random arrangement of our bundle. Units considered “good” by the Kilosort algorithm (consistently similar waveform shape and clean autocorrelations) were then manually curated and thrown out if their amplitudes or template presence drifted significantly over the course of the recording, or if cross-correlations with other units determined that they were duplicates. In the second case, the highest amplitude duplicate was kept, and the rest removed.

### Analyses and statistics

The following analyses were completed using custom MATLAB (R2022a) scripts, and the use of specific built-in MATLAB functions is noted when appropriate. These analyses were performed with the goal of testing for differences between these subregional pseudopopulations.

### Z-scoring and comparison of Z-scored firing rate across subregions

After spike sorting, we created three pseudopopulations by combining all neurons within each of the PFC subregions across all recording sessions. The first analysis involved Z-scoring the firing rate of every neuron around key behavioral events. This was done by summing the spikes in every 100 ms time bin five seconds before and after the sample, delay, or choice pokes for every correct trial (one hundred total bins for each poke). We then normalized each trial of each neuron across the time bins to create time-based Z-scores. These Z-scores were averaged across all correct trials in that neuron’s session to get the mean Z-score for every neuron around important DNMTP task events, and this result is depicted in the heatmaps seen in Figs. 2*B*, 3*B* and 4*B*. We could then take the mean across all neurons in each pseudopopulation to see how the regions differed in terms of simple firing rate changes over time (Figs. 2*C*, 3*C* and 4*C*).

To evaluate if the pseudopopulation Z-scored firing rate differed between subregions, we ran a separate one-way ANOVA (MATLAB function *anovan*) at every relevant time bin shown in the figures (this number was different between task phases). After uncorrected p-values for each ANOVA were found, we adjusted them for false discovery rate (FDR) using the Benjamini-Hochberg method ^65^. Each time point that still had a corrected p-value of < .05 was taken, and unpaired t-tests in that bin were conducted on each combination of subregional comparisons. The p-values from these multiple comparisons were then Bonferroni post-hoc corrected, and only comparisons with adjusted p-values still below .05 were considered significantly different at the corresponding timepoint. The above ANOVA strategy was also used to calculate significant subregional differences in the changes in Z-scored firing rate on incorrect vs correct trials in Fig. 8, except that that incorrect scores were subtracted from correct scores before analysis.

We took a related approach to analyze the percentages of neurons in each region that exhibited an increase or decrease by 0.5 Z-units (Figs. 2*D*, 3*D* and 4*D*). However, instead of running ANOVAs on each time point, we ran a χ^2^ test for homogeneity of proportions across the three groups (MATLAB function *crosstab*). Like the ANOVA approach, we also adjusted p-values for false discovery rate over time using the Benjamini-Hochberg method. If adjusted p-value was still less than .05 for a time bin, we ran separate pairwise χ^2^ tests for each combination of subregions and corrected for these multiple comparisons using the Bonferroni-Holm method ^66^. Any adjusted p-value below .05 after these conservative corrections was considered to represent a significant difference in the proportion of neurons either increasing or decreasing their firing rate between two groups at that time point. Moreover, this χ^2^ strategy was similarly employed in Fig. 8, to study the differences in percentages of neurons that increase or decrease their firing rates in response to incorrect trials.

### Determining retrospective sample location selectivity using permutation testing

Retrospective sample location selectivity was analyzed in several ways throughout the paper. In Figs. 2F, 3*F* and 4*F*, we used a permutation testing method to compare the raw spike counts in 100 ms time bins between left and right sample location trials (correct trials only). Since sessions rarely had an equal number of correct left and right sample trials, for each session we randomly subsampled trials from the greater of the two to match the number of trials from the lesser. Next, we randomly sampled two trials from a combined subpopulation of left and right samples, such that the spike counts for these random two trials could be from two left trials, two right trials, or a left and a right trial. We took the difference between two randomly sampled trials 1000 times and created a shuffled distribution of differences. We then calculated the true *mean* difference between all left and right trials, compared that to the shuffled distribution, and counted the number of shuffled differences that were greater than or less than the true mean difference. Neurons were considered to be significantly selective for a sample location if < 25 out of 1000 shuffled differences were greater in magnitude than the actual difference (approximating a p-value of < .025). This was done for every time point around key poke events.

### Support vector machines (SVMs) for cross-temporal decoding of sample location and prediction of poke context

To track the stability of location selectivity across time in each region, we trained linear SVMs (MATLAB *fitcsvm*) to decode retrospective sample location every 100 ms using binned raw spike counts from each neuron in a pseudopopulation on all correct trials. We then assessed how well each time point’s trained model could predict sample location based on the pseudopopulation activity at all time points (including the one it was trained on). A stable population representation would display above-chance predictive accuracy (> 50%) at time points far from the training time. A leave-two-out cross-validation scheme was applied to protect against overfitting the model. This consisted of holding out one trial from both left and right location samples and testing how the model trained on the remaining trials predicted the identity of these held out trials. Importantly, we subsampled all left and right trial counts to 12, which was the lowest left or right sample location trial count across all sessions, although most sessions had much more than 12. Our leave-two-out strategy was therefore repeated 12 times, with each value from this subsampling appearing once without replacement in the testing set. The prediction accuracy of the model for each subsample was calculated as the number of correct classifications of all held out trial combinations, out of 24. The overall subsampling procedure was repeated 20 times, for a total of 480 model predictions to calculate prediction accuracy for each cross-temporal comparison.

A second SVM was used in Fig. 5 to classify the poke context (sample, delay, or choice poke) in a brief window around the three poke events across correct trials. Because there were three contexts, a multiclass SVM was needed. We used the *fitcecoc* MATLAB function with a ‘one-vs-one’ coding design and 5-fold cross validation per comparison. We also ran a shuffled version of this model to confirm that the model was not overfit, and that chance level was ∼33%.

### General linear modeling of multiple task parameters over time

We used a general linear model (GLM, MATLAB function *fitlm*) to study how neurons in each region encoded multiple DNMTP task variables simultaneously around all task pokes. For Fig. 5, the GLM predictor variables included poke context (is the poke occurring during the sample, delay, or choice phase), sample location identity (right or left), choice location identity (left, center, or right), and outcome (correct or incorrect). This particular analysis with poke context (Fig. 5) was done in a brief time frame around poke events, because mice took inconsistent amounts of time to progress to different pokes, and we wanted to make sure the task phases in question were temporally well isolated. We took a similar GLM approach to Akam et al. (2021), in which a GLM was run on every time bin for every neuron centered around poke events, which initially included the contributions of all predictor variables (known as the “full” model). In other words, our GLM is attempting to predict the firing rate of each neuron at every categorical level (e.g. sample, delay, or choice poke) of our predictor variables at all time points in question.

The sum of squares error (SSE) of this full model represents the amount of residual variance in a neuron’s firing rate that cannot be explained by changes in the predictor variables. The full model should account for more firing rate variability and therefore have a relatively low SSE since it contains more relevant predictor variables. To uncover the extent to which pseudopopulations encoded singular task variables, we found the coefficient of partial determination (CPD) for each individual variable. This can be calculated by removing the singular task variables from the full model and running the GLM again on the reduced model. Since the reduced model should have a higher error term than the full, subtracting the full model from each reduced model should approximate the contribution of each removed variable to the full model’s firing rate prediction. The resulting number is the CPD for that singular variable, and it represents the percentage of total firing rate variability for a given neuron explained by that variable. A CPD was generated for each neuron at each time point, and the mean CPD for each pseudopopulation was reported over time and analyzed for significance with the same ANOVA approach as mentioned in *Z-scoring and comparison of Z-scored firing rate across subregions.* Using CPDs from all predictor variables, we could then compare the contributions of each variable to the *explainable* (not total) firing rate variance in each region.

For Fig 6., significantly strong CPDs (now measured across all relevant time points) for each variable were determined by comparing the true CPD value to a shuffled distribution of CPDs generated from randomly shuffling the relationship between firing rate and predictor variables 1000 times. As a very conservative cutoff, CPDs were only considered significant if *none* of the shuffled CPD values were higher than the unshuffled one, and we calculated the proportion of neurons at every time point that fulfilled this criterion. We then could determine differences in proportions of complex DNMTP task variable encoding between regions using the same χ^2^ approach employed at the end of the section *Z-scoring and comparison of Z-scored firing rate across subregions*. We then took these significant neurons and followed their sample location selectivity throughout the entire behavioral trial. Venn diagrams in Fig. 7 depict the percent of neurons with significant CPDs in each region taken from 3 bins (600 ms) on either side the labeled poke.

Among the most informative GLM outputs are beta-weights representing the strength and direction of the relationship between the firing rate of every neuron and the levels of each predictor variable. As another approach to visualize the stability of retrospective sample port location encoding, we recovered the maximum beta-weight amplitudes for every neuron with significant selectivity in a 600 ms time window around pokes. In this case, the beta-weights of significant neurons could be positive or negative with respect to the sample port location predictor variable, with negative signifying neurons that fired more on the left side of the box and vice versa. We quantified how these beta-weights changed over time using Pearson correlations (*r*, MATLAB function *corr*) comparing the significant beta-weight population vector at one point to the beta-weight population of those same neurons in the future. The significance levels of these correlations were also reported in Fig 7*C* and represent the confidence that the reported correlations are different from zero.

## Acknowledgements

We would like to thank Bakr Alkarawi for his help assembling the electrode implants used in this study. This work was financially supported by the Portland VA Research Foundation (PVARF), the OHSU Physician-Scientist Program to A.A., and the National Institute on Alcohol Abuse and Alcoholism (T32 AA007468 to A.S.).

## Contributions

A.S., A.A. and L.B. designed research; A.S. and L.B. performed research; A.S., R.O., L.B, and R.M. analyzed data; A.S. wrote the paper; A.S., L.B., R.O., R.M. and A.A. edited the paper.

## Ethics Declaration

Competing interest: None of the authors have any competing interest to declare.

## Notes

### Competing Interest Statement

The authors have declared no competing interest.

